# Dorsomedial striatal *Bmal1* deletion in medium spiny neurons alters EPSCs and dendritic morphology after chronic exposure to alcohol in female mice

**DOI:** 10.64898/2026.10.03.753483

**Authors:** Mahgol Darvishmolla, Richard Courtemanche, Shimon Amir, C. Andrew Chapman

## Abstract

Circadian clock genes in the striatum modulate alcohol drinking behavior in female mice, but the underlying synaptic mechanisms remain unknown. Medium spiny neurons (MSNs) of the striatum express D1- or D2-type dopamine receptors, which are differentially modulated by alcohol. Here we used whole-cell patch-clamp recordings to examine how deletion of the clock gene Bmal1 affects spontaneous excitatory postsynaptic currents (sEPSCs) in electrophysiologically identified D1 and D2 neurons of the female mouse dorsomedial striatum (DMS). Prior intermittent access to alcohol increased sEPSC amplitude in putative D1 neurons of wildtype mice without affecting sEPSC frequency, and this increase was accompanied by greater dendritic branching and complexity in biocytin-filled D1 MSNs. Bmal1 knockout had the opposite effect in D1 neurons, reducing sEPSC amplitude in both alcohol-exposed and alcohol-naive animals. In D2 neurons, alcohol produced only a non-significant trend toward increased sEPSC amplitude in wildtype mice, but a significant increase in Bmal1 knockout mice. Thus, local disruption of Bmal1 appears to block the alcohol-induced enhancement of sEPSCs in D1 neurons while unmasking a similar enhancement in D2 neurons. These findings may help explain how Bmal1 deletion in the DMS reduces alcohol consumption in female mice.

## 1. INTRODUCTION

Circadian rhythms synchronize internal physiological functions with environmental cues (zeitgebers) and modulate reward-related behaviours (Luo & Aston-Jones, 2009). Disruption of the circadian clock is strongly implicated in alcohol-related behaviors, and alcohol consumption in turn alters circadian rhythmicity (Tamura et al., 2021). PER2, CLOCK, and BMAL1 are core components of the molecular clock that regulates endogenous daily rhythms (Cox & Takahashi, 2019) and are linked to alcohol drinking and vulnerability to alcohol use disorder (AUD) (Meyrel et al., 2020; Wasielewski & Holloway, 2001). Polymorphisms in Bmal1 and Per2 in humans correlate with altered patterns of alcohol consumption, supporting the idea that genetic variation in circadian regulators contributes to AUD risk (Kovanen et al., 2010; Partonen, 2015; Valenzuela et al., 2016). In rodents, global deletion of Per2 or Clock, or reduced Clock expression in the nucleus accumbens (NAc), increases alcohol intake (Gamsby et al., 2013; Kirkpatrick et al., 2009; Rizk et al., 2022; Spanagel et al., 2005), and alcohol exposure alters the cyclic expression of PER2, which may contribute to circadian disturbances in individuals with AUD (McCarthy et al., 2013).

The striatum, which is implicated in reward-related behaviours including alcohol seeking, exhibits circadian rhythmicity through local expression of clock genes (Zacharias et al., 2025). More than 95% of striatal neurons are GABAergic medium spiny neurons (MSNs) (Matamales et al., 2009), which primarily express D1 or D2 dopamine receptors: D1 MSNs form the direct striato-pallidal pathway, and D2 MSNs form the indirect striato-pallidal pathway via the external globus pallidus and subthalamic nucleus (Lanciego et al., 2012). The striatum is thus a central structure in reward-related behaviour (Cox & Witten, 2019) through which circadian rhythms and alcohol use may interact (de Zavalia et al., 2021; Herrera et al., 2026).

### 1.1. Effects of Alcohol on the Striatum

The dorsomedial striatum (DMS) regulates drug and alcohol consumption (Corbit et al., 2012), and alcohol exposure enhances excitatory synaptic transmission within the striatum. Intermittent access to alcohol increases miniature EPSC frequency in the DMS (Wang et al., 2015) and produces a postsynaptic enhancement of glutamatergic corticostriatal transmission by increasing the amplitude of AMPA- and NMDA receptor-mediated EPSCs (Ma et al., 2017). Systemic ethanol administration can also facilitate long-term potentiation of glutamatergic inputs from the prefrontal cortex (Wang et al., 2015), and alcohol consumption facilitates glutamatergic transmission selectively in D1 MSNs of the direct pathway (Cheng et al., 2017; Wang et al., 2015).

Morphologically, alcohol also induces structural plasticity in MSNs of the dorsal striatum (Cheng et al., 2018; Clabough et al., 2021; Wang et al., 2015), which is thought to contribute to alcohol-drinking behavior (Chen et al., 2011). Acute neonatal alcohol exposure increases expression of postsynaptic density 95 (PSD-95) protein in MSNs (Alkaas et al., 2025) and increases dendritic length and branching (Clabough et al., 2021). In adult mice, alcohol increases dendritic length in D1- but not D2-MSNs (Wang et al., 2015) and alters signalling mechanisms that regulate dendritic spine morphology (Laguesse et al., 2018). Alcohol exposure can therefore have lasting effects on excitatory synaptic transmission and dendritic morphology in MSNs that may contribute to the persistence of alcohol-seeking behaviour.

### 1.2. Circadian Rhythm influence on striatal synaptic Transmission and Dendritic Spines

Excitatory corticostriatal inputs to MSNs may also be affected by clock gene expression and by disruption of the local striatal circadian rhythm. In ClockΔ19 mutant mice, the amplitude of AMPA-mediated miniature EPSCs in MSNs, and the AMPA/NMDA ratio, is reduced across the light/dark cycle relative to wild-type neurons, and ClockΔ19 MSNs are more hyperpolarized and less excitable both day and night (Parekh et al., 2018). This suggests that disrupting the circadian clock reduces excitatory synaptic transmission and neuronal excitability in the striatum. Circadian disruption may also affect dendritic complexity and spine morphology: in a mouse model expressing human cortical CLOCK, neurons expressing CLOCK show more complex dendritic branching, higher spine density, and increased spontaneous EPSC frequency (Liu et al., 2025), whereas CLOCK knockout in human induced pluripotent stem cell-derived neurons reduces both dendritic complexity and presynaptic vesicle number. Similarly, CLOCK knockout in layer V cortical pyramidal neurons reduces spine density in apical dendrites and primary branches (Li et al., 2017). Together, these findings suggest that dendritic architecture and synaptic connectivity are under circadian regulation in cortical neurons, and that the local circadian clock within the striatum may similarly regulate genes involved in cytoskeletal remodeling.

### 1.3. Combined Effects of Circadian Disruption and Alcohol Exposure in the DMS

Both alcohol consumption (Wang et al., 2015) and the circadian rhythm (Parekh et al., 2018) have powerful effects on excitatory transmission in the striatum, and the DMS emerges as a key candidate region through which circadian disruption may influence alcohol-related behavior. The present study therefore assessed the effects of chronic alcohol exposure and circadian disruption on excitatory synaptic activity and neuronal morphology of MSNs within the DMS. Our previous work showed that selective Bmal1 deletion across both the dorsal and ventral striatum decreases alcohol consumption in females and increases it in males, reflecting a sex-specific effect of striatal Bmal1 deletion on voluntary alcohol consumption (de Zavalia et al., 2021). Bmal1 knockout restricted to the nucleus accumbens, however, increased alcohol consumption in both sexes (Herrera et al., 2026), indicating that reduced clock function specifically in the dorsal striatum drives the reduced alcohol consumption seen in females after Bmal1 knockout across the dorsal and ventral striatum. Our subsequent behavioural studies showed that Bmal1 knockout restricted to the dorsomedial striatum, but not the dorsolateral striatum, reduces alcohol consumption in females without affecting males (Darvishmolla et al., 2026). Thus, while selective knockout of Bmal1 in the nucleus accumbens increases intake in females (Herrera et al., 2026), both global striatal knockout (Herrera et al., 2026) and dorsomedial striatal knockout (Darvishmolla et al., 2026) reduce alcohol consumption in females, identifying the dorsomedial striatum as a critical site for this sex-dependent effect.

Accordingly, the present study used viral-mediated Bmal1 deletion in the DMS of female mice to examine excitatory synaptic transmission and neuronal morphology in MSNs, in order to identify neural alterations that might underlie the reduction in alcohol consumption previously observed in females. Whole-cell recordings were used to assess neuronal excitability and to record spontaneous EPSCs from putative D1 and D2 MSNs that were discriminated electrophysiologically (Gertler et al., 2008) in acute brain slices from wild-type and Bmal1 knockout mice with or without intermittent access to ethanol. Dendritic morphology and spine density of MSNs were assessed using confocal microscopy to identify morphological differences associated with ethanol exposure and disruption of the local circadian clock.

## 2. MATERIALS AND METHODS

2.1. Animals and Surgery

Male and female C57BL/6J mice (12–16 weeks old) carrying floxed alleles of Bmal1 (Jackson Laboratory, Bmal1 fl/fl, stock no. 007668) were used in this study. Mice were housed individually in standard plastic cages at 21 ± 1 °C and 65 ± 5% relative humidity, under a 12-h/12-h light-dark schedule, with free access to food and water. All procedures were approved by the Concordia University Animal Care Committee (certificate no. 30000256). Animals were randomly assigned to Bmal1-knockout and control groups prior to viral injection surgery.

To delete Bmal1 in the DMS, Bmal1-floxed mice underwent bilateral stereotactic injections into the DMS under isoflurane anesthesia, delivered with a low-flow vaporizer system (SomnoSuite, Kent Scientific, Torrington, CT, USA). Mice were induced with 3–4% isoflurane in an induction chamber, then mounted in a stereotaxic frame, with anesthesia maintained via nose cone at 1–2% (0.2–0.5 L/min). Carprofen (5 mg/kg body weight) was administered as an analgesic.

Adeno-associated virus (AAV) expressing either Cre-recombinase and enhanced green fluorescent protein (EGFP) (AAV2/5-hSyn-CRE-EGFP, 1.0 × 10^12 gc/ml, Molecular Tool Platform, Laval University, Quebec, Canada) or EGFP alone (AAV2/5-hSyn-EGFP, 1.0 × 10^12 vg/ml, Molecular Tool Platform) was delivered to the DMS at the following coordinates: AP +0.98 mm, ML ±1.25 mm, DV −2.85 mm from bregma/skull surface, at an injection rate of 400 nl/min. The human synapsin (hSyn) promoter restricts transgene expression to neurons, limiting Bmal1 deletion to neuronal populations in the DMS without affecting glial cells (Kügler et al., 2003); however, Bmal1 deletion could still affect interneurons and cholinergic neurons and thereby alter synaptic inputs to D1 and D2 MSNs. The effectiveness of this viral strategy in reducing Bmal1 expression has been demonstrated previously in our laboratory (Darvishmolla et al., 2026).

### 2.2. Intermittent Access to Ethanol

At least 21 days after surgery, animals were habituated for three days to two water-filled bottles with ball-bearing spouts in their home cage. Mice then received 24-h sessions of free-choice access to water or 15% (v/v) ethanol (Peregud et al., 2021), every other day for 7 sessions; bottle positions were alternated in each session to control for side preference.

### 2.3. Electrophysiological Recordings

Recordings were obtained from brain slices at least 20 days after virus delivery and 2–3 weeks of intermittent alcohol exposure. Mice were anaesthetized with isoflurane and decapitated, and brains were rapidly extracted and cooled in oxygenated (95% O_2_/5% CO_2_) high-sucrose artificial cerebrospinal fluid (ACSF, 4 °C) containing (in mM) 2 KCl, 1.25 NaH_2_PO_4_, 7 MgSO_4_, 26 NaHCO_3_, 250 sucrose, 10 dextrose, and 0.5 CaCl_2_. Coronal slices (300 μm) containing the striatum were cut using a vibratome (Leica VT1200), held for 30 min at 32 °C in normal ACSF (in mM: 124 NaCl, 5 KCl, 1.25 NaH_2_PO_4_, 2 MgSO_4_, 2 CaCl_2_, 26 NaHCO_3_, 10 dextrose), and then kept at 22–24 °C until recording. Slices were placed in a recording chamber perfused with normal ACSF at 1.5–2.0 mL/min and visualized with an upright fluorescence microscope using a 40× water-immersion objective, differential interference contrast optics (Olympus BX51WI), and an XM-10 monochrome camera (Olympus, CellSens V1.8).

Whole-cell patch-clamp electrodes were pulled from borosilicate glass (1.0 mm OD, 3–5 MΩ) and filled with an internal solution containing (in mM) 140 K-gluconate, 5 NaCl, 2 MgCl_2_, 10 HEPES, 0.5 EGTA, 2 ATP-tris, 0.4 GTP-tris, and 0.1% biocytin (Sigma-Aldrich), pH adjusted to 7.25 with KOH (270–280 mOsm). Recordings were obtained only from neurons with soma diameter < 30 µm, to exclude cholinergic neurons and inhibitory interneurons, which have larger somata than medium spiny neurons (Chang & Kitai, 1985; Meredith et al., 1999); viral infection was confirmed by EGFP fluorescence before recording. Electrodes were lowered onto neurons in the DMS, tight seals were obtained (1.5–5 GΩ), and cells were allowed to stabilize for 10 min after achieving the whole-cell configuration.

Recordings were obtained with a Multiclamp 700B amplifier (Molecular Devices, San Jose, CA, USA), digitized (Digidata 1440A, Molecular Devices), and stored with pClamp 10.3 software (Molecular Devices). Access resistance averaged 46.7 ± 2.8 MΩ, and all cells had a resting membrane potential below −65 mV. The electroresponsiveness of MSNs was evaluated using 500-ms hyperpolarizing and depolarizing current steps from −200 to 100 pA in 20-pA increments from resting potential. Peak input resistance was measured at the largest voltage change during the −200 pA pulse, and steady-state input resistance was assessed just before the end of the current step. Action potential properties were assessed from the first spike evoked by positive current injection, and the number of spikes evoked at each current step was counted.

Voltage-clamp recordings of EPSCs were obtained at least 10 min after bath application of picrotoxin (50 µM, Abcam, Toronto, ON, Canada) and CGP55845 (1 µM, Tocris Bioscience, Bio-Techne Canada, Toronto, ON, Canada) to block GABA_A_ and GABA_B_ receptor-mediated inhibition. Cells were held at −80 mV with series resistance uncompensated, and spontaneous EPSCs were recorded for 5 min. Peak amplitude and decay time constant (tau) of EPSCs were analyzed using pClamp 10.8 software, and EPSC frequency was determined from the number of events detected over the 5-min recording. Effects of intermittent alcohol exposure and Bmal1 knockout were assessed using Holm-Sidak comparisons.

To allow intracellular diffusion of biocytin, the whole-cell configuration was maintained for 40–60 min, after which the electrode was slowly retracted from the soma. Apparent cellular capacitance typically decreased and access resistance increased on retraction, consistent with the soma remaining intact. Slices were then fixed in 4% paraformaldehyde at room temperature on a rotating shaker for 24 h.

### 2.4. Immunohistochemistry and Analysis of Dendritic Morphology

The morphology of biocytin-filled neurons was characterized using fluorescence imaging (Tan et al., 2023). Fixed slices were rinsed three times in PBS (0.2 M, 5 min each), then incubated in 0.1% Triton-X/PBS at 4 °C for 24 h. Slices were rinsed with PBS for 5 min and incubated in Streptavidin Alexa Fluor 568 (Thermo Fisher; 1:1000 in 0.3% Triton-X100/PBS) at 4 °C for 24 h. After three further PBS washes, slices were mounted on coverslips.

Micrographs were captured using a Zeiss Axio Observer 7 inverted microscope equipped with a Crest Optics Cicero spinning-disk module (Roma, Italy), a Hamamatsu Flash4.0 LT sCMOS camera (6.45 µm pixel pitch), a 60× Plan Apo objective (NA = 1.4), and an 89 North LDI5 laser launch with a 555 nm laser and a TXR (600/50m) emission filter. Image stacks were acquired at 0.2 µm z-spacing and 0.102 µm/pixel, and converted to two-dimensional images by maximum-intensity projection across the full stack in ImageJ (NIH, USA); no additional filtering or deconvolution was applied before projection.

Dendritic morphology was analyzed by Sholl analysis using the ImageJ Simple Neurite Tracer plugin, yielding measures of total dendritic length, number of branches, and number of branching points. Dendritic spine density was quantified on secondary and tertiary branches (Dobi et al., 2011), 50–100 µm from the soma and at least 20–30 µm from terminal tips (Wang et al., 2015). Images used for spine-density analysis were deconvolved with AutoQuant X3, and spines were counted per 10-µm segment (Mohan et al., 2019). All analyses were performed blind to treatment group, and data are expressed as mean ± SEM. Data were analyzed with one- or two-way ANOVA followed by Holm-Sidak post hoc tests.

## 3. Results

### 3.1. Identifying Putative D1- and D2-MSNs

Fluorescence imaging of EGFP during electrophysiological recordings confirmed efficient viral expression in the DMS (Fig. 1A), with large numbers of intensely fluorescent neurons distributed throughout the DMS in animals injected with either virus. Recordings were obtained from strongly fluorescent neurons near the slice surface (Fig. 1B, C). A total of 77 neurons were recorded; four were excluded because their firing patterns were consistent with fast-spiking GABAergic interneurons rather than medium spiny neurons.

**Figure 1.**
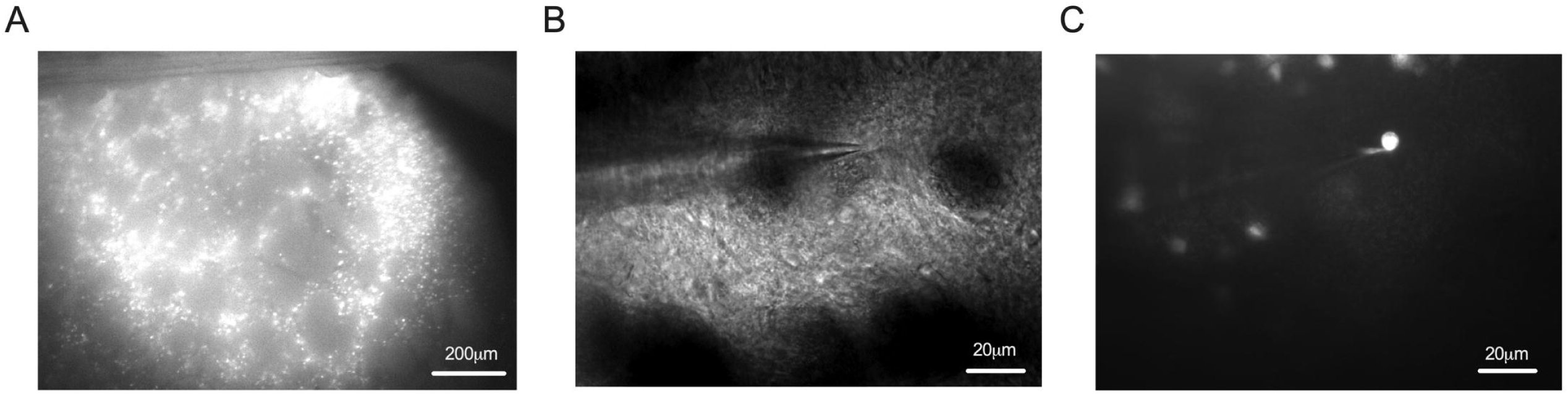
**A**: The image shows fluorescence from enhanced green fluorescent protein (E-GFP) near the injection site in the dorsomedial striatum, indicating Cre-mediated Bmal1 deletion. **B**: A brightfield image obtained during whole-cell recordings from a medium spiny neuron. The recorded cell is visible at the tip of the recording electrode. **C**: A fluorescence image of the same neuron shown in B.

D1 and D2 MSNs in the DMS differ in their intrinsic properties, including excitability, rheobase, and input resistance: D2 MSNs have higher input resistance and lower rheobase than D1 neurons (Gertler et al., 2008; Wang et al., 2015b; Willett et al., 2019a). Putative D1 and D2 neurons were therefore identified electrophysiologically using these criteria. After recording was complete, neurons with input resistance below 70 MΩ were categorized as putative D1-MSNs (mean 53.48 ± 3.51 MΩ, n = 39), and those above 70 MΩ as putative D2-MSNs (mean 115.80 ± 8.13 MΩ, n = 44). Rheobase current was also considered: neurons requiring 200 pA or more were classified as D1 (mean 260 ± 13 pA, n = 39), and those requiring less than 200 pA as D2 (mean 60 ± 15 pA, n = 44). These differences in input resistance and firing pattern between putative D1 and D2 neurons are illustrated in Fig. 2A, B.

**Figure 2:**
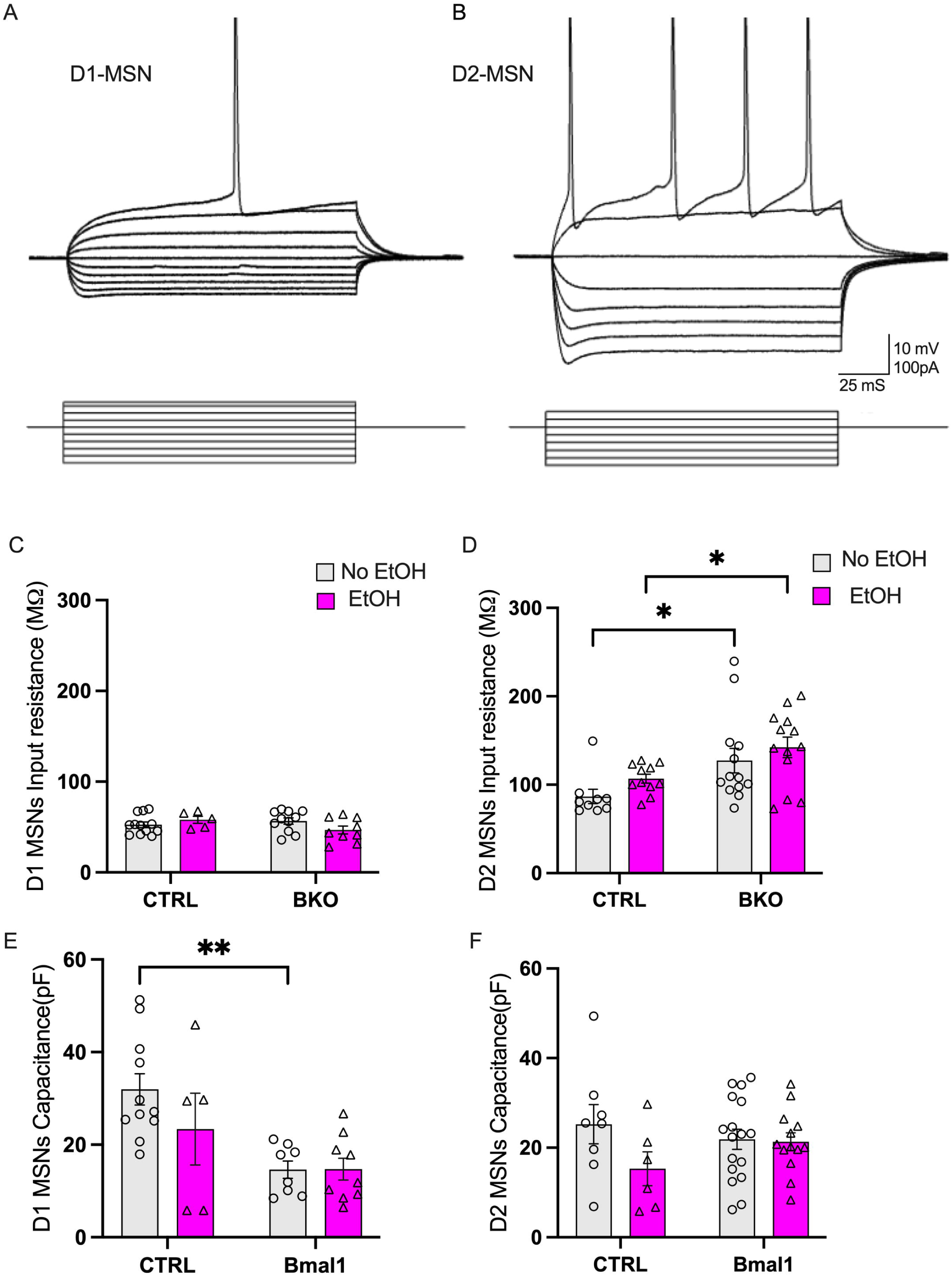
Bmal1 deletion, but not alcohol exposure, alters the intrinsic membrane properties of D1- and D2-MSNs. A, B: Representative membrane-voltage responses to hyperpolarizing and depolarizing current steps recorded from D1-MSNs (A) and D2-MSNs (B). C, D: Input resistance (Rin) of D1-MSNs (C) and D2-MSNs (D). Bmal1 deletion (BKO) significantly increased input resistance in D2-MSNs in both alcohol-exposed (EtOH) and non-exposed (No EtOH) groups (D), without affecting input resistance in D1-MSNs (C). Alcohol exposure had no significant effect in either cell type. E, F: Membrane capacitance (Cm) of D1-MSNs (E) and D2-MSNs (F). Bmal1 deletion in animals with no alcohol exposure significantly decreased capacitance in D1-MSNs but had no effect in D2-MSNs. Alcohol exposure did not significantly affect capacitance in either cell type. Data are presented as mean ± SEM; each point represents one neuron. Statistical comparisons were made using Holm-Šídák’s tests. *p < 0.05, **p < 0.01.

### 3.2. Bmal1 Deletion Selectively Affects Intrinsic Membrane Properties in D1- and D2-MSNs

We next asked whether Bmal1 deletion and/or alcohol exposure affect the intrinsic properties of D1- and D2-MSNs, which strongly influence neuronal excitability. In D1 neurons, input resistance was similar across all four groups: neither Bmal1 deletion (t(33) = 0.86, P = 0.395) nor alcohol exposure (t(33) = 0.90, P = 0.372) significantly altered input resistance relative to control (Fig. 2C).

In D2 neurons, alcohol exposure had no significant effect on input resistance in control animals (t(42) = 1.20, P = 0.236). Bmal1 deletion, however, increased input resistance in D2 neurons, both in alcohol-naive animals (t(42) = 2.49, P = 0.016) and in animals with intermittent alcohol exposure (t(42) = 2.302, P = 0.026) (Figure 2D), consistent with a Bmal1 knockout-induced enhancement of neuronal excitability. Thus, Bmal1 knockout increased input resistance in D2, but not D1, MSNs, regardless of alcohol exposure.

Membrane capacitance correlates with cell-membrane surface area and tends to be higher in larger neurons with more complex dendritic arbours (Taylor, 2012). In D2 neurons, capacitance did not differ significantly between groups, whether or not animals were exposed to alcohol (Bmal1 deletion: t(40) = 0.840, P = 0.40; alcohol: t(40) = 1.31, P = 0.197) (Fig. 2F). In D1 neurons, however, Bmal1 deletion decreased capacitance relative to control in alcohol-naive animals (t(29) = 3.638, P = 0.001), but this effect was not significant in alcohol-exposed animals (t(29) = 1.55, P = 0.130) (Fig. 2E), suggesting that Bmal1 knockout reduces D1 MSN membrane surface area and that this effect is attenuated by intermittent alcohol exposure.

### 3.3. Bmal1 Deletion Decreases sEPSC Amplitude in D1- but Not D2-MSNs

We analyzed the amplitude and frequency of spontaneous EPSCs to determine whether Bmal1 deletion and/or alcohol exposure affect excitatory synaptic inputs to DMS MSNs, and whether any changes occur pre- or postsynaptically. In wildtype D1 neurons, alcohol significantly increased sEPSC amplitude relative to alcohol-naive controls (t(34) = 2.51, P = 0.016). Bmal1 deletion was associated with reduced sEPSC amplitude both in alcohol-naive (t(34) = 2.07, P = 0.046) and alcohol-exposed animals (t(34) = 4.01, P = 0.0003) (Figure 3B), while neither alcohol nor Bmal1 deletion affected sEPSC frequency or decay time constant in D1-MSNs (Fig. 3C, D). These results suggest a postsynaptic facilitation of excitatory input to D1 MSNs by alcohol, which is counteracted by Bmal1 knockout.

**Figure 3:**
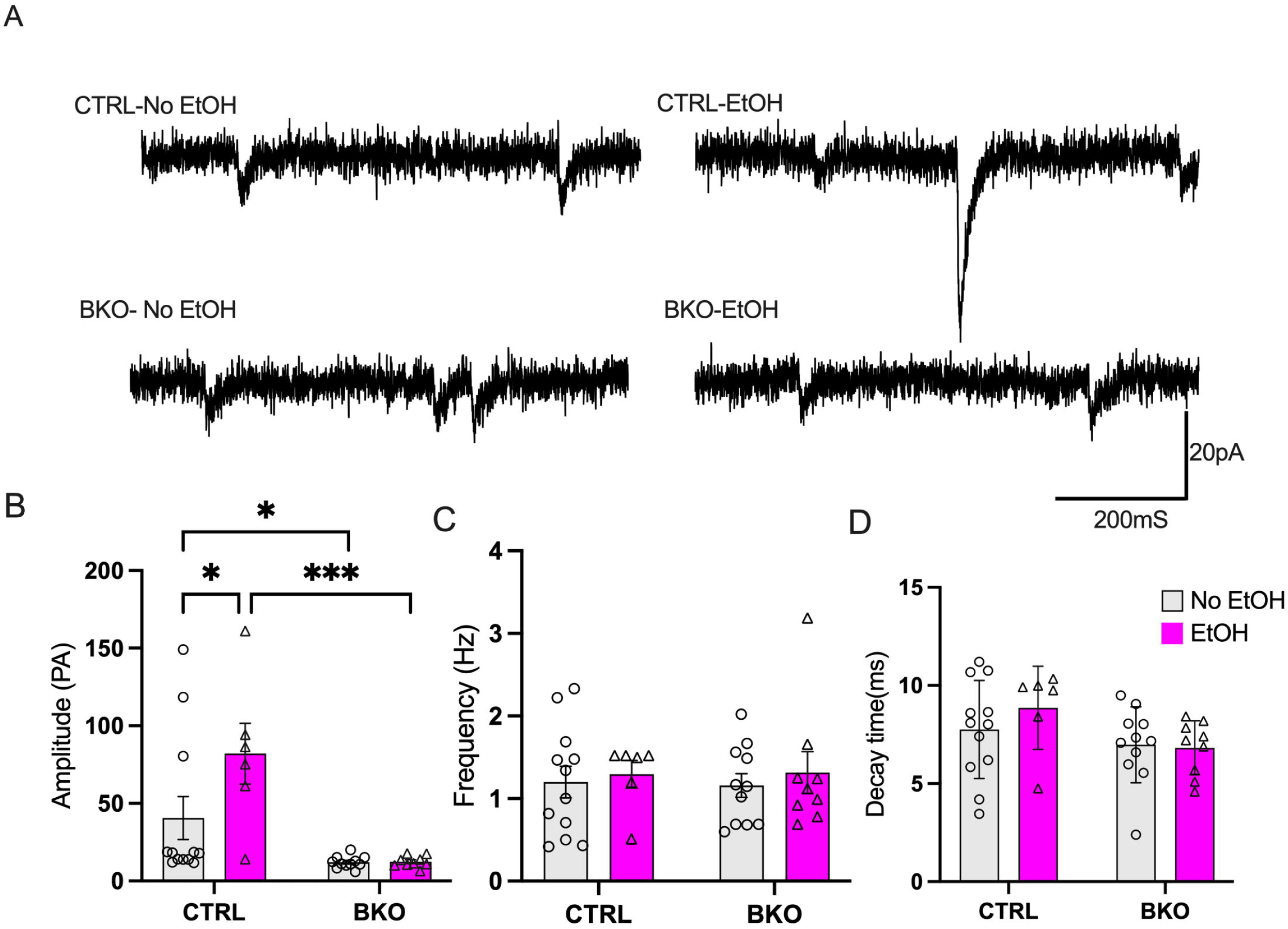
Bmal1 deletion decreases the amplitude of sEPSCs in D1 neurons. (**A**) Traces of the sEPSCs recorded from D1 neurons in four experimental groups. (**B–D**) Properties of sEPSCs in D1-MSNs in the DMS show the peak amplitude (**B**), frequency (**C**), and decay time constant (**D**). Alcohol exposure increased the sEPSC amplitude compared to the control group without Bmal1 deletion. Bmal1 deletion significantly reduced sEPSC amplitude in D1-MSNs compared to the control animals without Bmal1 deletion, and alcohol exposure further decreased the amplitude in Bmal1-deleted D1-MSNs; neither factor affected sEPSC frequency or decay time. (**B–D**). Calibration: 20 pA,100 ms.

In contrast, D2 neurons from wildtype animals showed no significant alcohol-dependent increase in sEPSC amplitude (t(40) = 0.97, P = 0.336), and Bmal1 deletion alone did not significantly affect sEPSC amplitude in alcohol-naive animals (t(40) = 0.22, P = 0.825). In alcohol-exposed animals, however, Bmal1 knockout was associated with significantly larger sEPSC amplitudes, both relative to wildtype alcohol-exposed neurons (t(40) = 3.50, P = 0.001) and relative to knockout animals without alcohol exposure (t(40) = 3.50, P = 0.001) (Fig. 4B). Neither alcohol exposure nor Bmal1 deletion affected sEPSC frequency or decay time in D2-MSNs (Fig. 4C, D). Together, these results indicate that the combination of Bmal1 knockout and intermittent alcohol exposure produces a postsynaptic facilitation of excitatory input to D2 MSNs.

**Figure 4:**
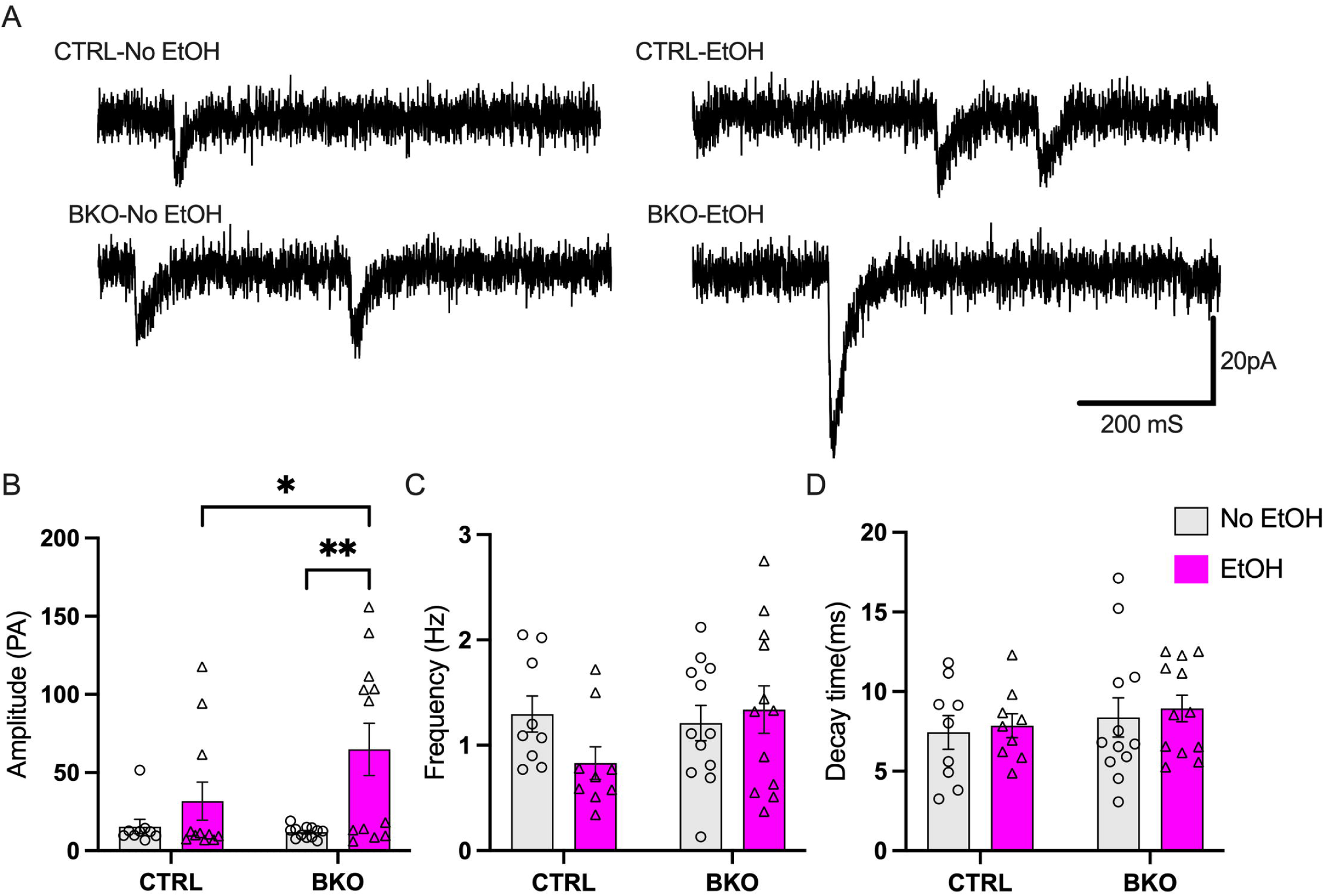
Bmal1 deletion increases the amplitude of sEPSCs in alcohol exposed D2 neurons. (**A**) Traces of the sEPSCs recorded from D2 neurons in four experimental groups. (**B–D**) Properties of sEPSCs in D2-MSNs indicate the amplitude (**B**), frequency (**C**), and decay time (**D**). Alcohol exposure increased the sEPSC amplitude in Bmal1-deleted D2-MSNs compared to Bmal1-expressing D2 neurons, and alcohol exposure to the Bmal1-deleted D2 neurons enhanced sEPSC amplitude relative to the Bmal1-deleted neurons with no exposure to ethanol; Bmal1 deletion alone did not alter amplitude, and neither factor affected sEPSC frequency or decay time. Calibration: 20 pA,100 ms.

### 3.4. Bmal1 Deletion Reduces Dendritic Complexity in D1-MSNs of the DMS

To identify a possible structural basis for the effects of Bmal1 deletion on intrinsic properties and sEPSC amplitude, we analyzed dendritic branching and complexity in D1- and D2-MSNs using a Sholl analysis, which quantifies dendritic complexity by counting intersections between dendrites across concentric circles of increasing radius centered on the soma (Binley et al., 2014). Neuron images were flattened for analysis (Fig. 4A, B, G, H).

In D1 neurons, the number of Sholl intersections varied significantly with distance from the soma across groups (distance × group interaction: F(36,240) = 7.175, P < 0.0001), indicating that the effects of alcohol exposure and Bmal1 deletion on dendritic complexity depended on distance from the soma. As shown in Figure 4C, alcohol increased the number of intersections at 40–100 μm from the soma relative to alcohol-naive controls (P < 0.0001, marked by *), whereas Bmal1 deletion reduced intersections at 60–70 μm relative to control (P < 0.0001, marked by +). The combination of alcohol exposure and Bmal1 deletion reduced dendritic complexity at 60–110 μm relative to alcohol-exposed controls (P = 0.001, marked by #). Together, these results show that alcohol exposure increased the number of intersections in wildtype neurons, an effect that was absent in Bmal1 knockout neurons.

Similarly, Bmal1-deficient D1 neurons had shorter total dendritic length than Bmal1-expressing neurons, both in alcohol-naive (t(20) = 3.04, P = 0.006) and alcohol-exposed animals (t(20) = 2.641, P = 0.015) (Fig. 4D), whereas alcohol exposure alone had no significant effect on total dendritic length (t(20) = 0.67, P = 0.509).

Dendritic complexity was also assessed from the total number of branches and branching points (Fig. 4E, F). Bmal1-deficient neurons had fewer total branches than controls, both in alcohol-naive (t(20) = 2.70, P = 0.013) and alcohol-exposed animals (t(20) = 3.72, P = 0.001), whereas alcohol exposure alone did not significantly affect branch number in either wildtype (t(20) = 0.45, P = 0.651) or Bmal1 knockout animals (t(20) = 0.56, P = 0.577) (Fig. 4E).

The number of branching points showed a different pattern: alcohol exposure increased branching points relative to alcohol-naive controls (t(20) = 4.00, P = 0.0007), but not in Bmal1 knockout animals (t(20) = 0.33, P = 0.724), and Bmal1 deletion alone did not significantly alter branching points relative to control (t(20) = 1.03, P = 0.314) (Fig. 4F).

### 3.5. Bmal1 Deletion Does Not Affect Dendritic Complexity in D2-MSNs of the DMS

Sholl analysis of D2 MSNs showed a broadly similar number of intersections across groups (Fig. 4I), although a significant distance × group interaction (F(36,192) = 1.55, P = 0.031) reflected a small but significant reduction in intersections at 70 μm from the soma in alcohol-exposed Bmal1-deficient neurons relative to alcohol-naive controls (P = 0.026, marked by #). Neither Bmal1 deletion (t(21) = 0.04, P = 0.962) nor alcohol exposure (t(21) = 0.15, P =

0.883) significantly affected total dendritic length relative to control (Fig. 4J). Total branch number was likewise unaffected by Bmal1 deletion (t(24) = 0.69, P = 0.494) (Fig. 4K), and neither alcohol exposure (t(24) = 1.36, P = 0.183) nor Bmal1 deletion (t(24) = 1.99, P = 0.581) significantly affected the number of branching points relative to control (Fig. 4L).

### 3.6. Bmal1 Deletion Decreases Spine Density on D1-MSNs in the DMS

Because Bmal1 deletion reduced sEPSC amplitude in D1-MSNs but not D2-MSNs, we asked whether this was accompanied by corresponding changes in dendritic spine density. Fig. 5A shows representative images of dendritic segments from D1 MSNs in each of the four groups. Bmal1 knockout was associated with significantly reduced spine density both in alcohol-naive (t(24) = 4.46, P = 0.0002) and alcohol-exposed animals (t(24) = 2.75, P = 0.011) (Fig. 5B), whereas alcohol exposure alone had no significant effect on spine density in either wildtype (t(24) = 0.57, P = 0.572) or Bmal1 knockout animals (t(24) = 1.14, P = 0.263).

**Figure 5:**
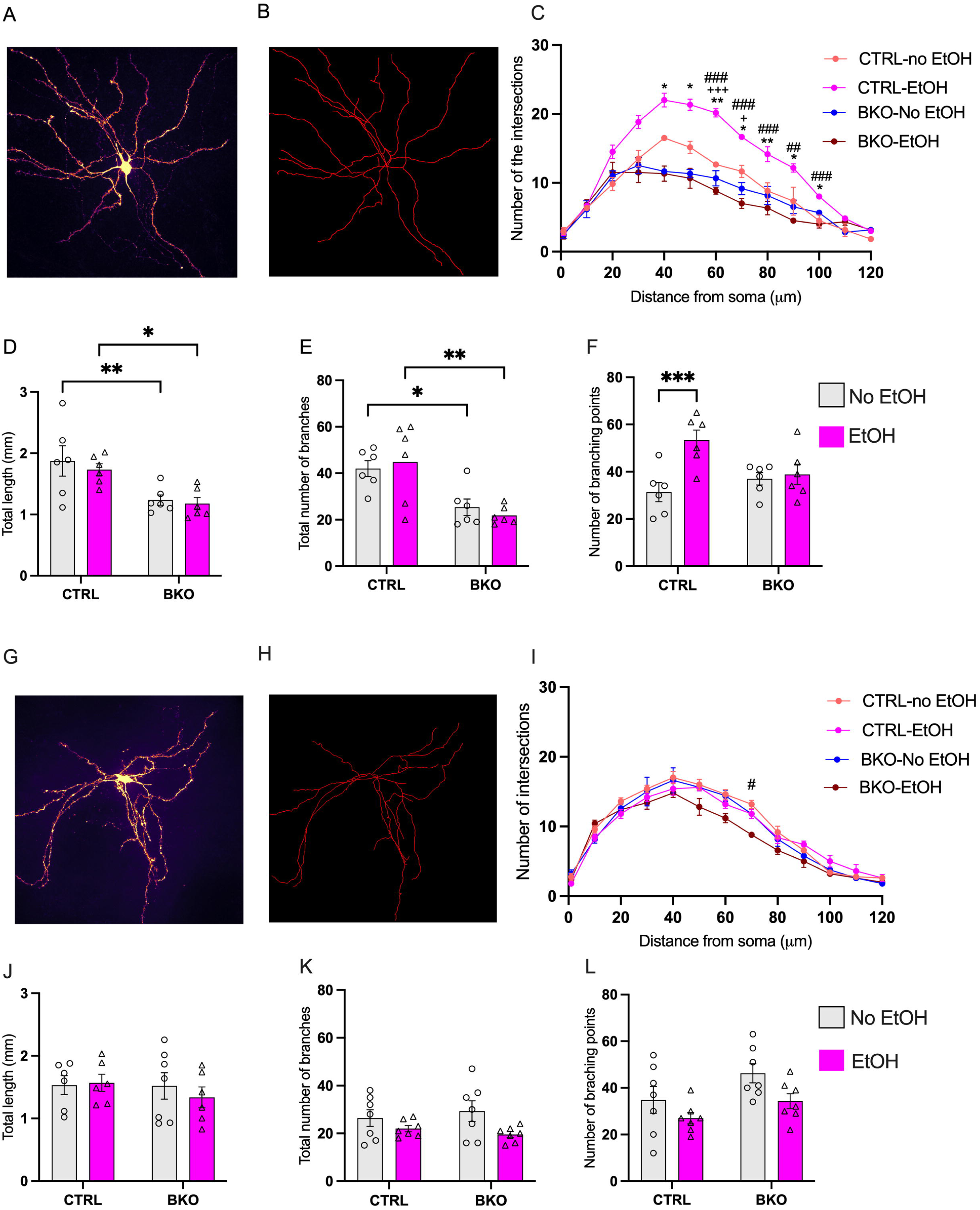
Bmal1 deletion reduces dendritic complexity in D1- but not D2-MSNs of the DMS. **A, G**: Representative fluorescence images of a D1-MSN (A) and D2-MSN (G) and their dendritic reconstructions (B & H). **C:** Sholl analysis of D1-MSNs shows the number of dendritic intersections with concentric circles as a function of radial distance from the soma. A significant distance × group interaction was detected in which alcohol exposure increased intersections at 30–100 μm from the soma compared to the wildtype group without alcohol exposure (*), and *Bmal1* deletion reduced intersections at 50–60 μm (□) relative to the wildtype cells without ethanol exposure. There was a genotype × alcohol interaction in which complexity was reduced at 50–100 μm (#) compared to the wildtype with ethanol. **D-F**: Dendritic morphology of D1-MSNs. *Bmal1* deletion significantly reduced total dendritic length (D) and total number of branches (E) compared to the wildtype neuron without alcohol exposure. Alcohol exposure decreased the total length and the total number of branches in Bmal1-deleted D1 neurons relative to the wildtype group with alcohol exposure (D&E). Alcohol significantly increased the total number of branching points compared to the control group without *Bmal1* deletion (F). **I:** Sholl analysis of D2-MSNs indicated that alcohol exposure reduces the number of intersections at 70 μm from the soma in *Bmal1*-deleted D2 neurons relative to wildtype neurons (#). J–L: Neither *Bmal1* deletion, alcohol exposure, nor their interaction affected any of these measures in D2-MSNs. Sholl profiles (C, I) were analyzed by two-way repeated-measures ANOVA (distance × group); morphometric measures by two-way ANOVA (genotype × alcohol). All were followed by Holm-Sidak’s test. In the Sholl panels: *p < 0.05 (alcohol vs. control), □p < 0.05 (*Bmal1* deletion vs. control), #p < 0.05 (genotype × alcohol interaction).

**Figure 6:**
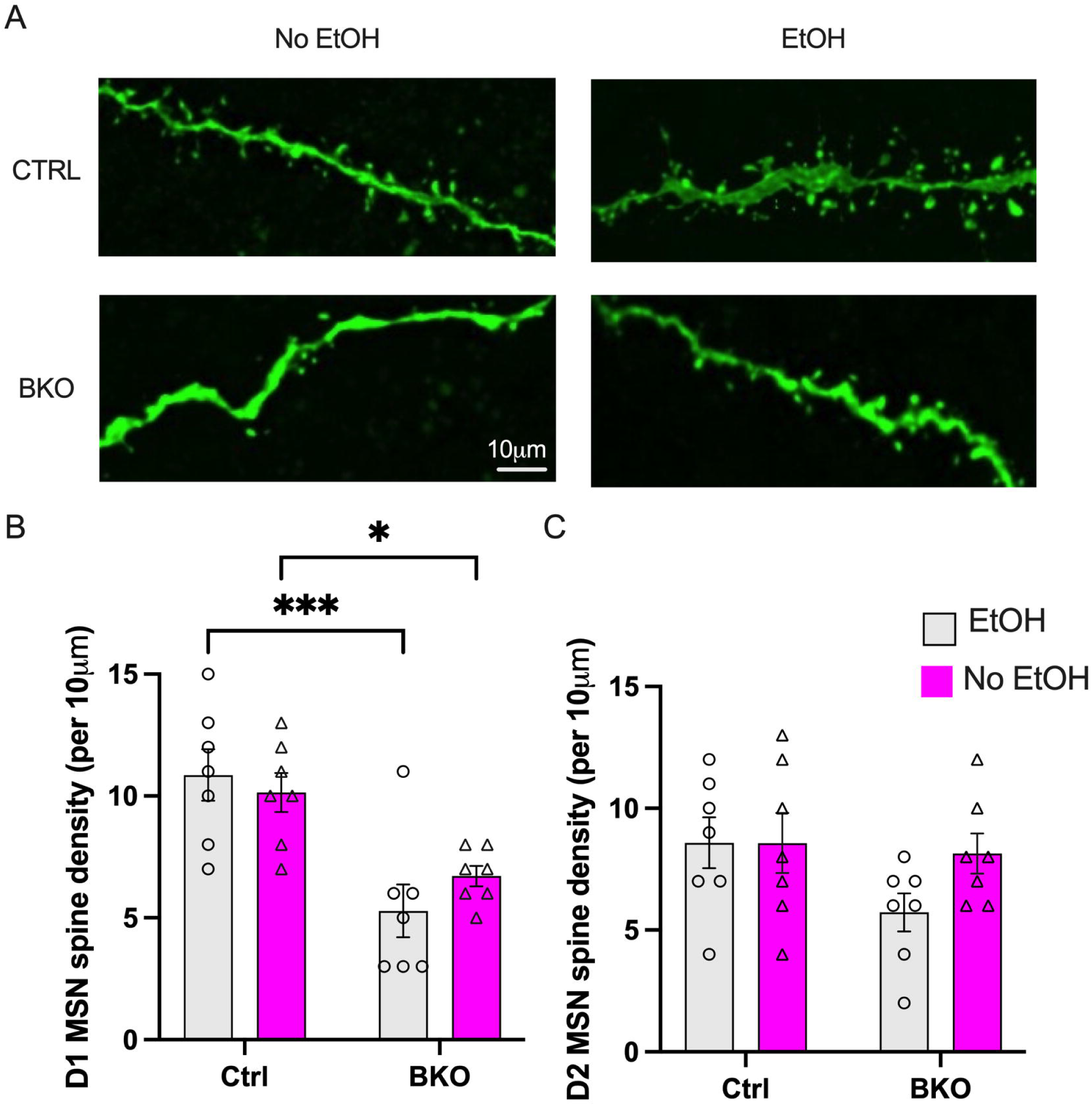
Bmal1 deletion decreases dendritic spine density on D1- but not D2-MSNs in the DMS. A: Representative confocal images of segments of D1-MSN dendrites in each of the four experimental groups (CTRL–no EtOH, CTRL–EtOH, BKO–no EtOH, BKO–EtOH). B: Quantification of dendritic spine density (spines/10 μm) in D1-MSNs. Bmal1 deletion decreased spine density relative to the wildtype groups without or with alcohol exposure. C: Quantification of spine density on D2-MSNs; neither Bmal1 deletion nor alcohol exposure significantly altered spine density.

In D2 neurons, alcohol exposure had no effect on spine density in either wildtype (t(24) = 0, P > 0.99) or Bmal1 knockout animals (t(24) = 0.30, P = 0.761), and Bmal1 deletion alone did not significantly alter spine density relative to control (t(24) = 2.04, P = 0.518) (Fig. 5C). Dendritic spine density is therefore strongly and selectively reduced by Bmal1 knockout in D1 MSNs, but not D2 MSNs, and is not markedly affected by alcohol exposure in either cell type.

## 4. Discussion

This study shows that Bmal1 deletion and chronic alcohol exposure differentially modulate the electrophysiological and morphological properties of D1- and D2-MSNs in the dorsomedial striatum. Alcohol exposure did not affect input resistance or firing properties of MSNs, but did significantly increase sEPSC amplitude in D1 neurons of wildtype mice, along with an increase in branching points and dendritic intersections in Sholl analysis. Bmal1 knockout had the opposite effect in D1 neurons, reducing sEPSC amplitude in both alcohol-exposed and alcohol-naive animals, so that the alcohol-induced facilitation of sEPSCs was observed only in wildtype mice. Bmal1 deletion also shaped the effect of alcohol in D2 MSNs: alcohol produced only a non-significant trend toward larger sEPSCs in wildtype mice, but a marked increase in mean amplitude in Bmal1 knockout mice. Thus, local Bmal1 disruption appears to block the alcohol-induced enhancement of sEPSCs in D1 neurons while facilitating a comparable enhancement in D2 neurons. These electrophysiological and anatomical changes underscore a cell type-specific role for Bmal1 in the DMS, and may contribute to the cellular mechanisms underlying the reduction in alcohol consumption previously observed in female mice after DMS Bmal1 deletion (Darvishmolla et al., 2026).

D1- and D2-MSNs were identified from their intrinsic electrophysiological properties as established in the literature (Gertler et al., 2008; Willett et al., 2019b) rather than with transgenic reporter lines or cell-type-specific viruses, which would provide more definitive classification. However, the distinct effects of alcohol exposure and Bmal1 knockout on the two putative cell populations suggest that this classification was reliable.

Alcohol consumption is a goal-directed behavior that involves the DMS (Corbit et al., 2012; Yin & Knowlton, 2006). We found that intermittent alcohol exposure increased sEPSC amplitude in DMS D1-MSNs of wildtype mice, and in D2-MSNs of Bmal1 knockout mice. Alcohol exposure has previously been shown to enhance excitatory synaptic activity in DMS neurons (Ma et al., 2017), and inhibiting glutamatergic transmission in the DMS suppresses alcohol-drinking behavior (Wang et al., 2012). Our finding that sEPSC facilitation in wildtype animals occurred in D1 but not D2 neurons is consistent with previous reports of alcohol-induced enhancement of excitatory cortical input specifically to D1 MSNs. Alcohol self-administration enhances NMDA receptor activity in D1-MSNs, and inhibiting D1 neurons and their cortical inputs reduces alcohol consumption (Roltsch Hellard et al., 2019). Alcohol potentiates both AMPA (Wang et al., 2015) and NMDA receptor-mediated responses (Cheng et al., 2017) in D1-MSNs but not D2-MSNs (Cheng et al., 2017; Wang et al., 2015). Presynaptic cortical terminals onto DMS

MSNs can carry both D1 and D2 receptors (Lu et al., 2021; Paspalas & Goldman-Rakic, 2005), and alcohol exposure has notably been found to facilitate large-amplitude responses mediated by D2-expressing terminals that contact D1 MSNs (Lu et al., 2019). Together with prior reports that alcohol enhances both AMPA- and NMDA receptor-mediated glutamatergic transmission in D1-MSNs (Roltsch Hellard et al., 2019; Wang et al., 2015), our finding that alcohol increases sEPSC amplitude in DMS D1-MSNs further supports the idea that D1-MSNs are a primary postsynaptic target of alcohol-induced glutamatergic potentiation.

In contrast to the alcohol-induced facilitation of sEPSCs, Bmal1 deletion powerfully reduced sEPSC amplitude in D1 MSNs regardless of alcohol exposure. Because amplitude rather than frequency was reduced, Bmal1 likely regulates postsynaptic receptor sensitivity, density, or function rather than presynaptic excitability (Gonzalez-Islas & Wenner, 2006). This finding aligns with previous work on how circadian clock disruption affects reward circuitry: ClockΔ19 mutant mice show reduced AMPA receptor-mediated miniature EPSC amplitude in the nucleus accumbens (NAc) with no change in presynaptic release probability, attributable to diurnal reductions in trafficking and surface expression of the AMPA receptor subunit GluA1 (Parekh et al., 2018). Because Bmal1 dimerizes with Clock to drive tissue-specific transcription (Trott & Menet, 2018), the similar effects of Bmal1 and ClockΔ19 deletion on EPSC amplitude suggest that a functional Bmal1-Clock complex is normally required to maintain baseline AMPA receptor density at D1 MSN synapses. That Bmal1 deletion reduced sEPSC amplitude only in D1 neurons suggests that Bmal1 regulates postsynaptic glutamatergic strength specifically within D1 neurons that mediate the direct-pathway.

In contrast to the reduction in sEPSC amplitude we observed after Bmal1 knockout, knockdown of Neuronal PAS Domain Protein 2 (Npas2) in the NAc selectively enhances AMPA receptor-mediated mEPSC amplitude in D1-MSNs without affecting D2-MSNs (Parekh et al., 2019). These opposing effects may reflect functional differences between Bmal1-Npas2 and Bmal1-Clock heterodimers. Npas2 and Clock have overlapping roles but produce distinct transcriptional outputs depending on cellular context (DeBruyne et al., 2007; Landgraf et al., 2016): Npas2 can substitute for Clock in the suprachiasmatic nucleus (SCN), the primary circadian pacemaker, but peripheral tissues of Clock-deficient mice lack circadian rhythms, indicating that Clock is essential for circadian oscillation outside the SCN (Landgraf et al., 2016).

Npas2 is a transcriptional partner of Bmal1 (Reick et al., 2001) that is highly enriched in D1-MSNs relative to D2-MSNs (Ozburn et al., 2015), and the facilitation of EPSCs after Npas2 knockdown (Parekh et al., 2019) suggests that Npas2 normally suppresses excitatory transmission. Bmal1 knockout in our animals would prevent dimerization of Bmal1 with both Clock and Npas2, yet we did not observe the facilitation of EPSC amplitude seen after Npas2 knockdown in NAc neurons. This suggests that Npas2 may suppress excitatory transmission through mechanisms independent of its dimerization with Bmal1, though the extent to which Bmal1 and Npas2 interact to regulate synaptic strength in the DMS remains unclear. One possibility is that the Bmal1-Npas2 complex normally restrains transcriptional activation of AMPA receptor machinery: when Npas2 is deleted and the Bmal1-Clock complex predominates, AMPA receptor expression may increase, raising mEPSC amplitude. By contrast, Bmal1 deletion eliminates both arms of this regulatory axis, collapsing E-box-driven transcription and reducing postsynaptic glutamatergic strength in D1-MSNs.

Unlike in D1 neurons, mean EPSC amplitude in D2 neurons was increased by alcohol exposure, and this effect was significant only in Bmal1 knockout mice. Clock gene mutation in the ventral tegmental area (VTA) decreases expression of the GABA_A_ receptor α1 subunit, and the resulting reduction in inhibitory tone increases VTA neuron excitability (McClung et al., 2005). Reduced inhibitory tone in the DMS following Bmal1 knockout may similarly have increased D2 neuron excitability, contributing to the alcohol-induced enhancement of sEPSCs we observed. However, because alcohol also potentiates GABAergic transmission onto D2 neurons (Cheng et al., 2017), it remains unclear why this facilitation appeared only in alcohol-exposed knockout mice and not in wildtype mice.

The alcohol-induced enhancement of sEPSC amplitude in D1 neurons of wildtype animals was accompanied by increased complexity of dendritic arbour (Fig. 5A-F): alcohol exposure increased the number of branching points and intersections in D1 neurons without altering branch number or total dendritic length, and had no effect on dendritic complexity or length in D2 neurons. These findings are consistent with previous reports that alcohol increases dendritic intersections in D1, but not D2, neurons (Wang et al., 2015).

In contrast, the Bmal1-induced reduction in sEPSC amplitude in D1 neurons, seen in both alcohol-exposed and alcohol-naive groups, was associated with reduced intersections, total dendritic length, branch number, and spine density in D1 neurons (Figure 5A-F). Consistent with these findings, membrane capacitance proportional to neuronal surface area (Taylor, 2012), was reduced in D1 neurons from Bmal1 knockout versus wildtype animals, an effect that was significant in alcohol-naive animals. Clock knockout similarly reduces dendritic arborization in the mouse frontal cortex (Liu et al., 2025) and in human brain tissue (Li et al., 2017). Dendritic complexity in D2 neurons, by contrast, was largely stable, aside from a reduced number of intersections at 70 µm from the soma in alcohol-exposed Bmal1 knockout animals, indicating that the effects of Bmal1 deletion on dendritic morphology are highly cell type-specific.

Dendritic spine structure is primarily supported by actin, which mediates changes in spine morphology (Nithianandam & Chien, 2018; Honkura et al., 2008), and Bmal1 deletion in astrocytes has been shown to impair the actin cytoskeleton and alter the morphology of distal astrocytic processes (Ali et al., 2020). Together, our findings identify Bmal1 as a potential regulator of structural plasticity in D1-MSNs: the selective reduction in dendritic complexity, spine density, and membrane capacitance following Bmal1 deletion suggests that Bmal1 is required to maintain D1 MSN morphology, potentially through regulation of the actin cytoskeleton. These structural changes may in turn limit the number and efficacy of glutamatergic synapses in D1-MSNs, contributing to the reduced sEPSC amplitudes observed in Bmal1-deficient D1-MSNs.

## 5. Conclusion

In summary, alcohol exposure facilitates spontaneous synaptic activity in D1 neurons of wildtype animals, while Bmal1 deletion selectively reduces dendritic complexity, spine density, membrane capacitance, and excitatory synaptic strength in D1-MSNs, leaving D2-MSNs largely unaffected. Bmal1 deletion blocked the alcohol-induced facilitation of sEPSC amplitude in D1 neurons, and instead facilitated an alcohol-induced enhancement of excitatory synaptic strength in D2 neurons. Given that alcohol disrupts the corticostriatal pathway controlling goal-directed behavior (Renteria et al., 2018) and increases postsynaptic glutamatergic activity in D1-MSNs (Wang et al., 2015), and that clock genes contribute to glutamatergic transmission in MSNs (Parekh et al., 2018), our findings suggest that Bmal1 expression in D1 MSNs of the DMS maintains excitatory corticostriatal connectivity within the direct pathway of the striatum and supports alcohol reinforcement, whereas Bmal1 deletion instead facilitates alcohol-mediated effects on excitatory transmission in the indirect pathway. These findings provide candidate cellular mechanisms for the female-specific reduction in alcohol consumption previously observed following Bmal1 deletion in the DMS (Darvishmolla et al., 2026).

## Acknowledgment

We thank Dr. Chris Law for his invaluable technical support in the Center for Microscopy and Cellular Imaging (CMCI) at Concordia University.

## Author Contributions

Conceptualization, data curation: MD, RC, SA, CAC; supervision, validation, funding & project administration: RC, SA, CAC; methodology, data acquisition and analysis: MD, CAC; writing, original draft: MD, CAC; writing – review and editing: MD, SA, RC, CAC.

